# Individual variation in the accuracy of saccades to sounds is correlated with the properties of eye movement-related eardrum oscillations (EMREOs)

**DOI:** 10.64898/2026.05.13.724941

**Authors:** Jesse L. Herche, Cynthia D. King, Jennifer M. Groh

**Affiliations:** Dept. of Neurobiology, Dept of Psychology and Neuroscience Duke Institute for Brain Sciences Center for Cognitive Neuroscience Duke University, Durham, NC 27708

## Abstract

Calibration of sound localization behavior in species with mobile eyes requires not only accurate visual input but also accurate oculomotor signals across the lifespan. The recent discovery of eye movement-related eardrum oscillations suggest that oculomotor signals may be incorporated into auditory processing at the level of the ear. One inference of this discovery is that individual variation in such signals might be correlated with individual variation in sound localization accuracy. Here, we tested this hypothesis in humans with normal hearing. We discovered that there is considerable variation in the accuracy of saccades to sounds even in normal individuals: median horizontal errors ranged from 2-6°, and median vertical errors could be as large as 36°. We separated the subject pool into groups with “good” performance (median vectorial error < 8°) vs “poor” performance (median vectorial error > 10°) and evaluated their respective EMREOs. The EMREOs differed across the two groups in how horizontal saccade amplitude and vertical initial eye position were encoded, with considerable individual variation in the “poor” group. These results are consistent with the interpretation that EMREOs are associated with underlying processes that ensure the accuracy of sound localization.

**HIGHLIGHTS:**

- The accuracy of eye movements to look at sounds varied across individuals, with median errors spanning a greater than 10-fold range. This wide range is surprising given that the participants passed screening for normal hearing.
- “Good” vs “poor” sound localizers exhibited differences in their eye movement-related eardrum oscillations (EMREOs)
- EMREOs differed in both horizontal and vertical sensitivity, for both saccade amplitude (horizontal) and initial eye position (vertical), and the differences varied in timing with respect to saccade onset.
- We interpret the results under the theory that poor sound localization may be a consequence of poor eye movement encoding, which is essential for accurately linking visual and auditory space.

## INTRODUCTION

Eye movement-related signals have recently been discovered in the ears of humans and non-human primates (Gruters, Murphy et al. 2018, Bröhl and Kayser 2023, King, Lovich et al. 2023, Lovich, King et al. 2023, Lovich, King et al. 2023, Abbasi, King et al. 2025, Sotero Silva, Kayser et al. 2025, King and Groh 2026, King, Zhu et al. 2026, Leon, Ramos et al. 2026, Sotero Silva, Brohl et al. 2026). These eye movement-related eardrum oscillations (EMREOs) have been postulated to play a role in linking visual and auditory spatial information (Gruters, Murphy et al. 2018, Lovich, King et al. 2023). The theory behind this is that sound localization is based on cues that are anchored to the head and ears (interaural timing and level difference cues, as well as spectral cues), whereas visual spatial information derives from where on the retina the image lies. In species with mobile eyes, there is no fixed correspondence between retinal and head/ear centered cues: only by including information about the angle of gaze with respect to the head is it possible to discern whether a given visual object and a given sound arise from the same location in the physical world.

Correct binding between the visual and auditory scenes is important not just for processing the immediate sensory environment, but also over time for aiding in calibrating the accuracy of sound localization (for reviews, see (King 2009, Opoku-Baah, Schoenhaut et al. 2021)). In humans, head diameter and the consequent separation between the two ears roughly doubles from birth to adulthood, thus the mapping between particular values of interaural timing and level differences to physical locations in the external world must be continuously adjusted. Work in barn owls suggests that visual input provides that instructive signal (Knudsen and Knudsen 1985), but barn owls do not move their eyes, so an oculomotor signal is not required for ascertaining the correct binding between visual and auditory input in that species. In humans, incorporation of accurate eye movement/position signals is needed together with that visual input to calibrate the accuracy of sound localization cues.

It follows, then, that if someone localizes sound poorly, the problem might lie with the accuracy of information about eye movement/position. Specifically, failing to incorporate accurate information about eye movements/changes in eye position into the coordination between auditory and visual processing could lead to impairments in the accuracy of sound localization. Potentially, anomalies in the EMREO could lead to such impairments, and might be evident when examining individual variation in EMREOs in comparison to sound localization performance.

To test this hypothesis, we evaluated the accuracy of sound localization in N=23 human subjects using an auditory saccade task, and we investigated the correlation between their performance at localizing sounds and the properties of their EMREOs. We found that there is considerable intersubject variability in the accuracy of sound localization, with median horizontal errors ranging from 2-6 degrees, and median vertical errors as high as 36 degrees. Furthermore, subjects who were poor at localizing sounds (median vectorial error > 8 degrees) showed anomalies in their EMREOs: the dependence on horizontal saccade amplitude was generally weaker in this group than in the group that showed good sound localization performance. The temporal profile of the dependence on vertical initial eye position was also altered and included anomalous phase inversions across time. Such effects could indicate that anomalous eye-movement related corollary discharge signals are present in these subjects. Overall, these results support the interpretation that EMREOs reflect underlying mechanisms that are important for linking visual and auditory space, such that anomalies in this signal are associated with poorer sound localization performance.

## RESULTS

N=27 human subjects with normal hearing and corrected to normal vision were recruited for participation in these studies (see Methods). Sound localization performance and EMREOs were recorded in separate sessions in sound attenuation chambers lined with echo absorbing material. Eye movements were tracked using video eye trackers. Four of these subjects were excluded from further analysis due to an excess amount of blinking during the sound localization task, leaving a final dataset of N=23 subjects.

The sound localization sessions involved a task designed to assess accuracy in both the horizontal and vertical dimensions, with either visual or auditory stimuli presented as saccade targets (Figure 1A). Each trial began with an initial (visual) fixation position followed by a saccade target (visual or auditory). The initial fixation positions and saccade targets ranged in horizontal location and were vertically offset from one another as shown (fixation positions 8 ° below the target row). Accurate performance required making saccades to both the correct horizontal and vertical positions of the targets. The trials with visual targets also helped ensure the accurate calibration of the eye tracker. Visual stimuli consisted of dots projected onto an acoustically transparent screen concealing the locations of auditory speakers. Sounds consisted of broadband noise (55 dB SPL) presented from those speakers. All conditions were randomly interleaved.

**Figure 1.**
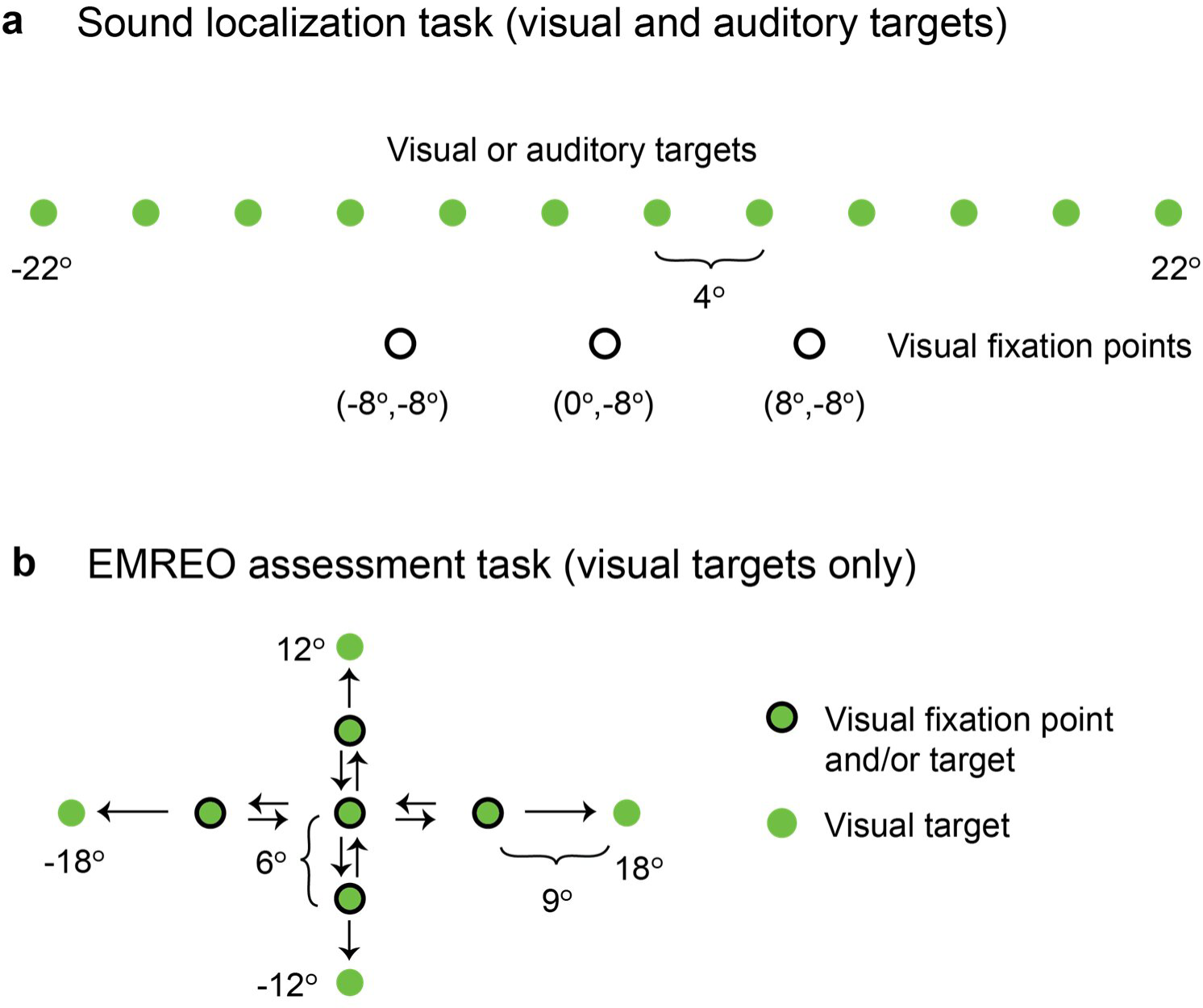
Sound localization and EMREO assessment tasks. **a**. The sound localization task involved an initial visual fixation stimulus from a row of three possible locations 8 ° below the horizontal meridian. After participants had fixated that stimulus for 400-600 ms, another target appeared, from a row of locations on the horizontal meridian. This second target was either visual, to ensure proper calibration of the eye tracking system, or auditory (broadband noise at 55 dB SPL), to allow assessment of sound localization accuracy. The visual stimuli consisted of white dots on a dark background and were presented by computer projector onto an acoustically transparent screen that prevented the participants from seeing the speakers at any time. **b**. The EMREO assessment task was conducted in a separate sound attenuation chamber to ensure quiet earbud microphone recordings. Participants fixated one of 5 possible locations (straight ahead, 9° left or right, or 6° up or down) before making saccades either left or right or up or down from those positions as indicated by the arrows. No diagonal fixation-target combinations were used. Visual stimuli were presented on a computer monitor. No sounds were presented in this task.

Participants varied in their performance. Figures 2 and 3 show the results for two sample participants, with the subject in Figure 2 (S104) showing better performance than the subject in Figure 3 (S120). Panels A-B show eye traces vs time for the visual (A) and auditory (B) trials, color coded according to the target location. For simplicity, only the horizontal dimension of eye position is shown, and only trials from the central fixation point are displayed.

**Figure 2.**
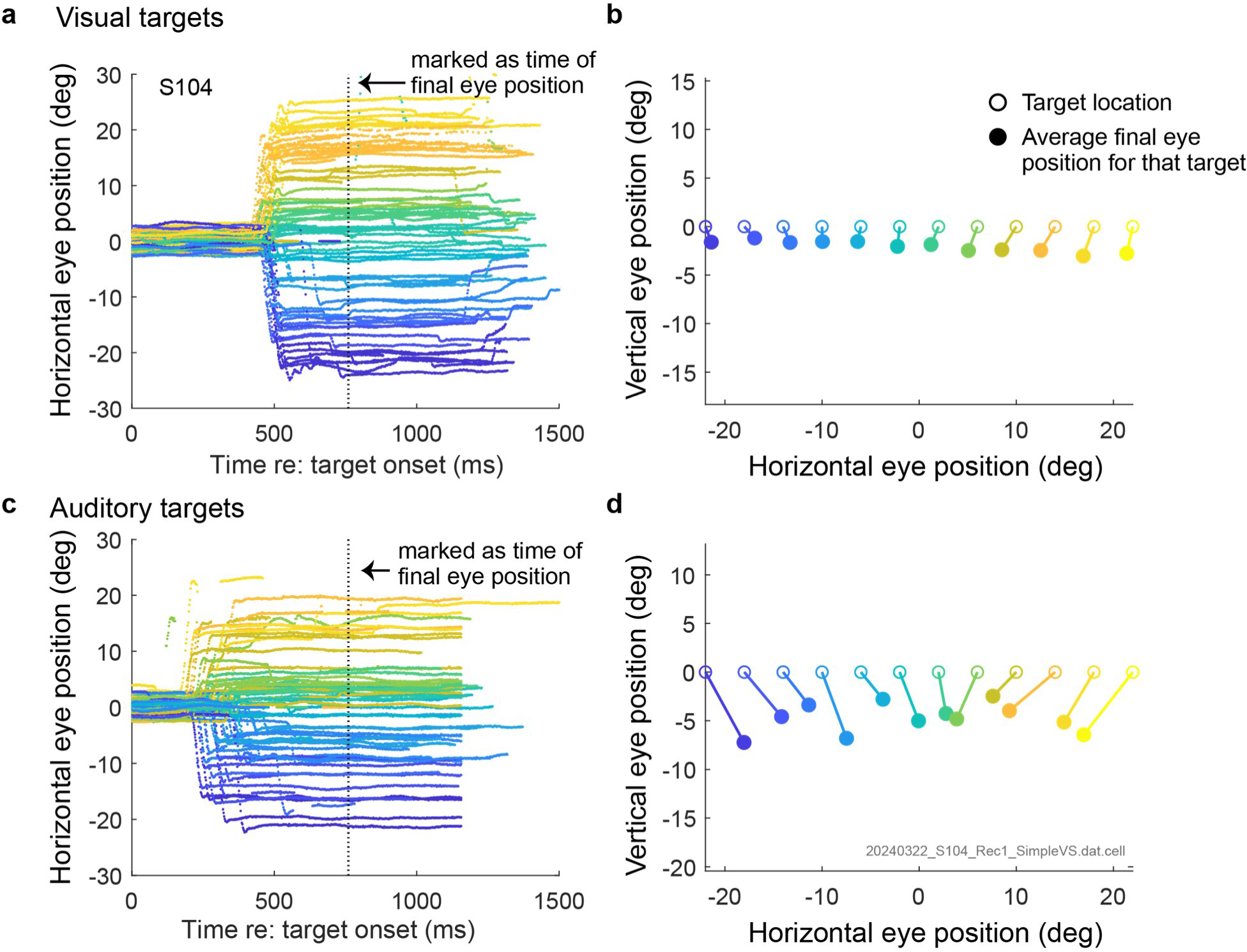
Results from a subject (S104) with “good” performance on the sound localization task. **a**. Horizontal eye position vs. time for visual trials from the central fixation position. The vertical dashed line indicates the point in time at which final eye position was assessed, as described in the main text. The color coding corresponds to the different target locations. **b**. Average final eye position, using the same color coding for target position (open circles) as in **a**. c, d. Same as for a, b but for auditory trials. This subject’s good sound localization performance is indicated by the fact that the average final eye positions “sort” well in the horizontal dimension, although the range is somewhat compressed relative to the actual range of targets, and the vertical error is modest.

**Figure 3.**
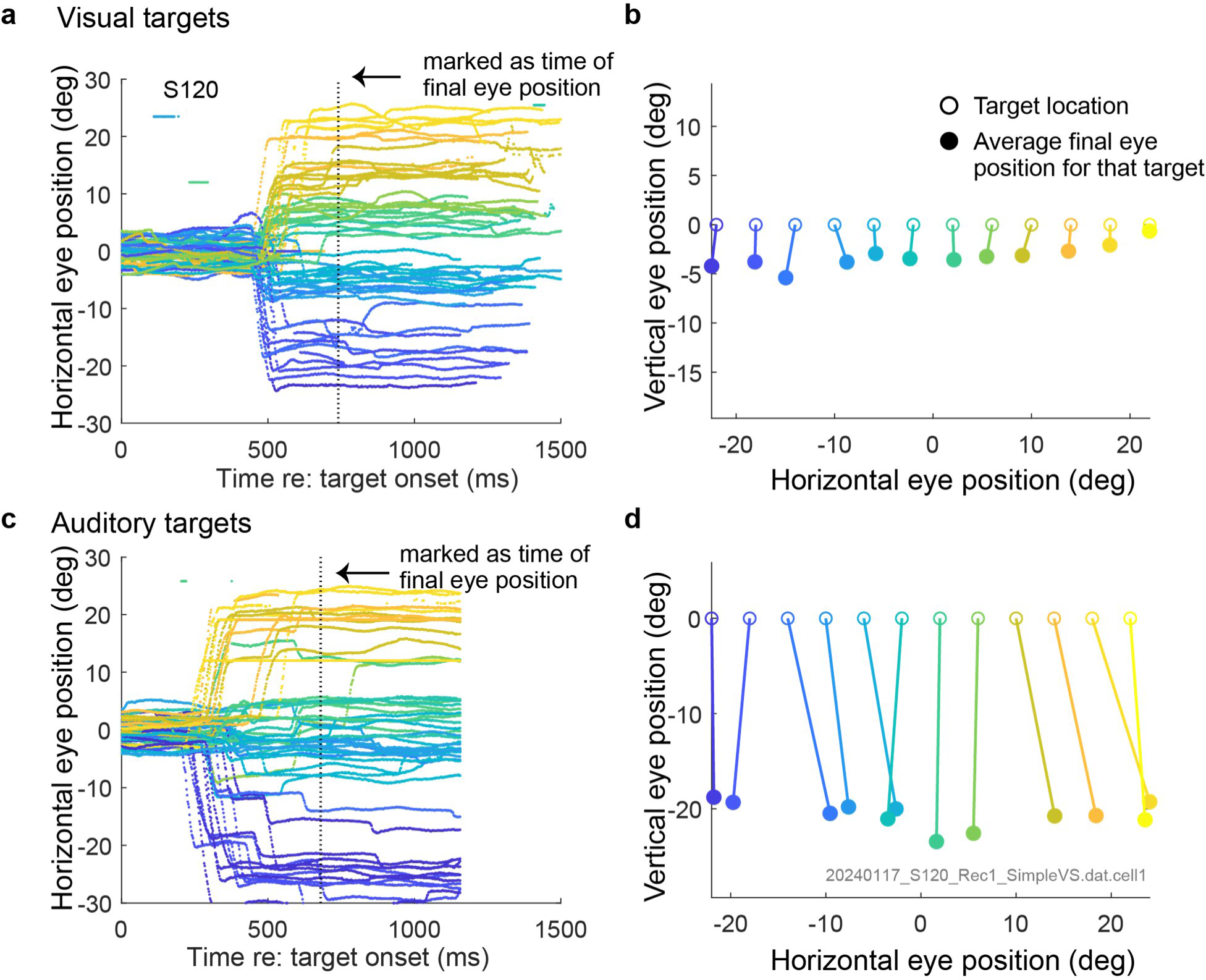
Results from a subject (S120) with “poor” performance on the sound localization task. Same format as Figure 2. This subject exhibited normal performance on visual trials (**a,b**), but greatly mislocalized the sounds in the vertical dimension ( **d**). Performance in the horizontal dimension also showed some inaccuracies, such as grouping different target locations together.

Both subjects performed well when making saccades to visual stimuli: the reaction times are consistent across trials and generally only one saccade was needed to arrive at the target. For S104 (Figure 2), this is true of auditory trials as well. But for S120 (Figure 3), many trials involved two saccades to arrive at the sound location. It is a known phenomenon that many subjects make two saccades to look at sounds (Jay and Sparks 1990).

Since our main interest is the “final” saccade endpoint, we assessed eye position at a fixed point in time after target onset as the subject’s best report of target position. We chose this time point separately for each subject and separately for visual and auditory trials as follows: we computed a regression of horizontal eye position as a function of horizontal target location, in 20 ms increments from 500 to 760 ms after target onset, and we chose the time point that had the largest product of the r^2^ and slope values – that is, a combination of low variance across trials and a good fit to the target location itself. The best fitting time point is shown for each subject and target modality in Figures 2 and 3 A and B with a vertical dotted line.

With a time point in hand, we could then compute the average eye position for each target and overall median error in both horizontal and vertical dimensions as shown in Figures 2 and 3 panels C and D. For visual trials (C and D), both subjects show good horizontal localization accuracy and a modest vertical offset. The overall median errors were 2.6 and 4.1 degrees, respectively.

For auditory trials, the errors were larger than for visual trials for S104 (Figure 2): an overall median of 6.4, due to a combination of a median horizontal error of 3.0 degrees and a median vertical error of 4.8 deg. These slightly larger errors compared to visual trials are comparable to those reported with highly experienced participants in the older saccadic sound localization literature (Zambarbieri, Schmid et al. 1982, Jay and Sparks 1990, Frens and Van Opstal 1995, Populin 2008). For S120, however, performance was much worse on auditory than for visual trials, specifically in the vertical dimension. The subject exhibited an overall median error of 21.6 degrees, based largely on a median vertical error of 20.8 degrees as well as a more modest median horizontal error of 3.6 deg.

The performance variation evident across these two participants was present across the population as a whole: Figure 4 shows the median vectorial (Figure 4A) and horizontal and vertical errors (Figure 4B) across the population. Based on gaps across these distributions, the population was subdivided into 2 groups: a “Good sound localizers” group, in which participants had a median horizontal error less than 3.5 degrees and vertical median error less than 6 degrees (dashed lines) (N=9 of 23) and a “ Poor sound localizers” list, that failed to meet these criteria (N=14 of 23). The largest vectorial median error in the “good” group was 7.3° (the minimum was 3.5°), whereas the smallest vectorial error in the “poor” group was 10.1° (the maximum was 36.2°).

**Figure 4.**
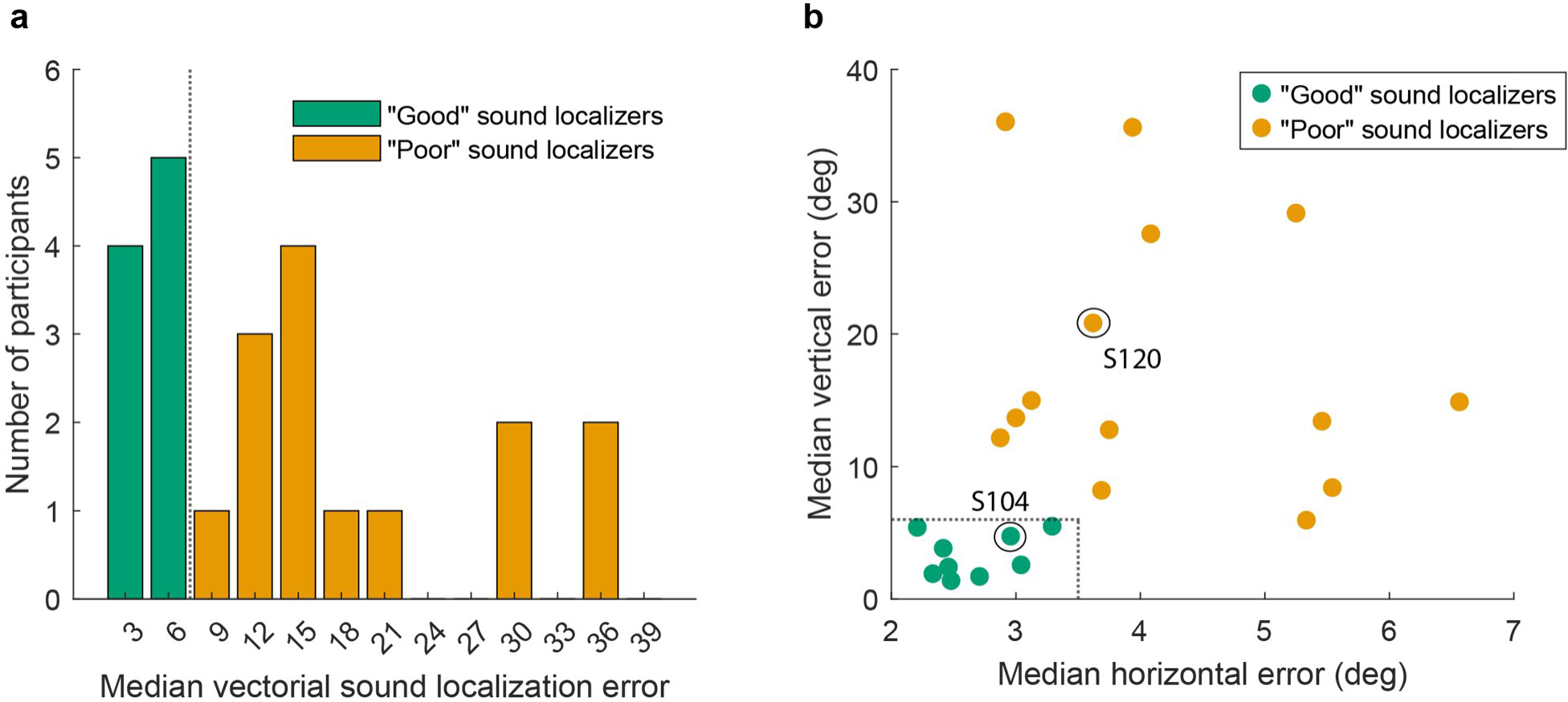
Median vectorial (a) and horizontal and vertical sound localization errors (b) in the subject pool. Note the different ranges of the X and Y axes in (b). The dashed lines indicate the criteria for inclusion in the “good” sound localizer group. The circles and subject numbers indicate the subjects illustrated in Figures 2 and 3.

We then evaluated the EMREOs in these different subject populations. As noted above, the EMREO sessions were conducted separately from the sound localization sessions, because EMREO measurement requires placement of earbuds in the ear and does not involve the delivery of any external sounds (but see (Sotero Silva, Brohl et al. 2026)). Rather, participants perform a visually guided saccade task, sampling both the horizontal and vertical dimensions as shown in Figure 1B.

Results are shown in Figure 5 for the two example subjects shown earlier in Figures 2 and 3 for a subset of the target locations and combined across the two ears. We have previously shown that when trying to read out the location of a saccade target from the EMREO recordings, it is beneficial to take the difference between the two ears for reading out the horizontal dimension and the average of the two ears for the vertical dimension (Lovich, King et al. 2023). Accordingly, for the horizontal dimension, Figure 5A shows the average microphone recordings for the left ear minus the right ear, as a function of whether the target was 9 degrees to the right (red) or 9 degrees to the left (blue). For the subject with good performance in the sound localization task (top), the traces show large-amplitude oscillations, well balanced across the two saccade directions, and cleanly out of phase with each other. For the subject with poorer performance (bottom), the oscillation is quite obviously present, but it is much smaller in amplitude and exhibits a temporal pattern that is atypical, with late deflections at a lower oscillatory frequency. For the vertical dimension (Figure 5B), results are shown for the average across the two ears, and are color coded for up (green) or down (purple). Again, the “good” subject shows a clean oscillatory signal, whereas the “poor” subject exhibits unusual temporal patterns and a smaller signal overall.

**Figure 5.**
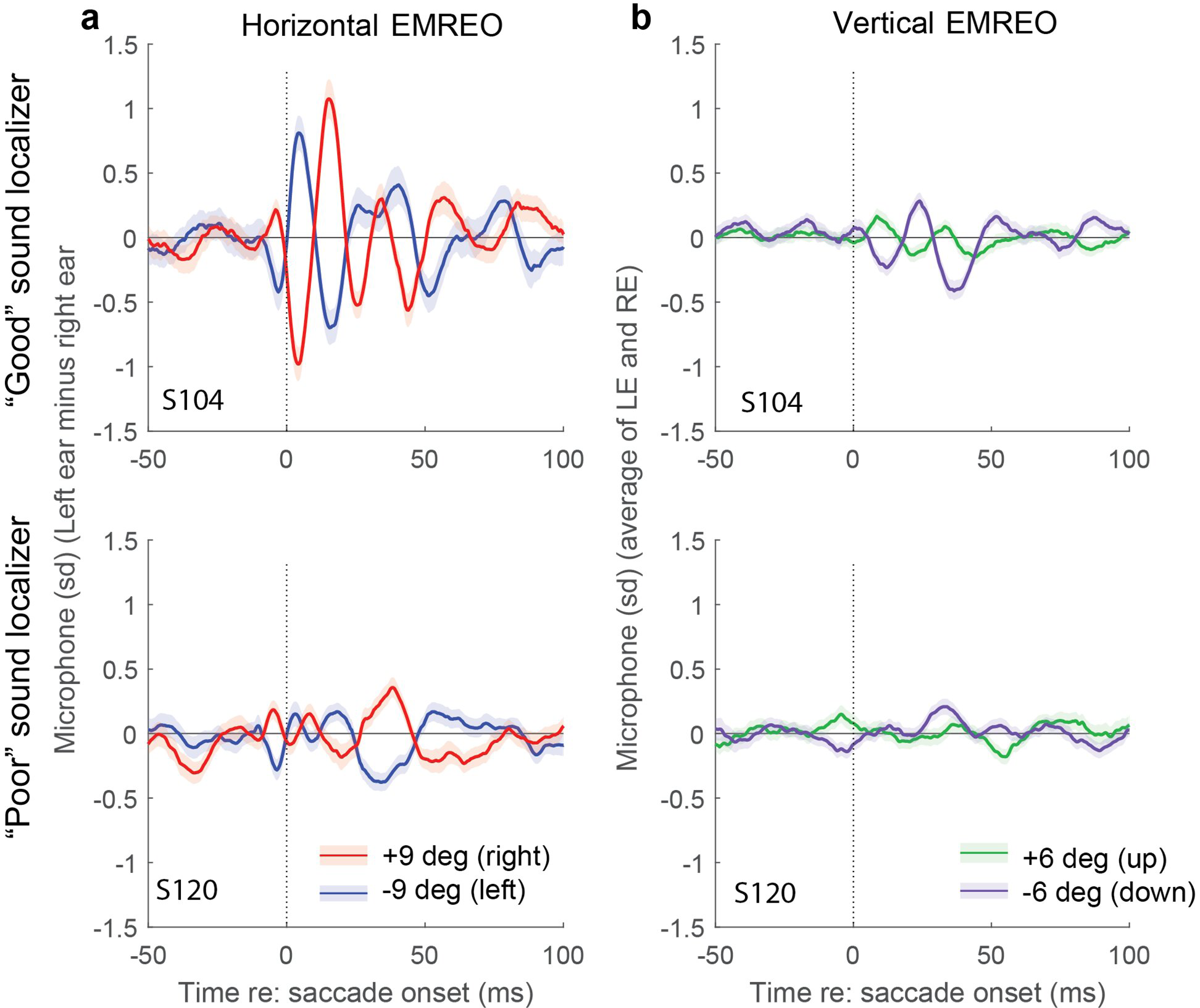
Horizontal and vertical EMREOs of the two example subjects illustrated in Figures 2 and 3. **a**. For the horizontal dimension, these plots show the signal recorded in the left ear minus the signal recorded in the right ear, for 9° rightward (red) and leftward (blue) saccades, pooled across initial fixation positions. Shading shows the standard error across trials. **b**. For the vertical dimension, the signals are averaged across the left and right ears as well as pooled across initial fixation positions; 6° upward saccades are shown in green and downward in purple. Taking the difference across the two ears has previously been shown to be beneficial for reading out the horizontal dimension and averaging is more beneficial for the vertical dimension (Lovich, King et al. 2023). Regardless of horizontal vs. vertical dimension, the signal (and the difference across saccade directions) is overall larger in S104 (top), who showed good performance on the sound localization task, then for S120 (bottom), who showed poor performance.

These patterns were borne out at the population level. To evaluate the results across ears, dimensions (horizontal and vertical), and as a function of both saccade amplitude and initial fixation position, we fit the microphone data with a regression equation as follows:

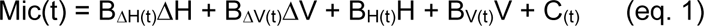

where H and V refer to the horizontal and vertical locations of the initial fixation position on a given trial; ΔH and ΔV refer to the target location relative to that initial fixation position, and the coefficients B_H(t)_ B_V(t)_ B_ΔH(t)_ B_ΔV(t)_ are time-varying coefficients designating the weighting of those factors. In addition, the regression has a time-varying intercept term C_(t)_ which typically captures asymmetries in the signal between leftward and rightward and/or upward and downward.

Figure 6 shows the population averages of these regression coefficients for both horizontal and vertical dimensions and for both saccade amplitude and initial eye position components. As in Figure 5, results are combined across ears differently for the horizontal and vertical dimensions: the right ear is subtracted from the left ear for assessing the information contained in the EMREO with respect to the horizontal dimension, and the two ears are averaged for assessing the signals in the vertical dimension. Shading indicates the standard error of the mean, and dots at the top of each panel indicate points in time for which a t-test indicated a statistically significant difference between the two groups at a p<0.05 level. At the lower left of each panel, the percentage of time points that were significant for the time window shown (−50 to 100 ms with respect to saccade onset) is provided. Cluster-based permutation testing (Maris and Oostenveld 2007) was then conducted to assess the overall significance of the difference between “good” and “poor” localizers for each regression component.

**Figure 6.**
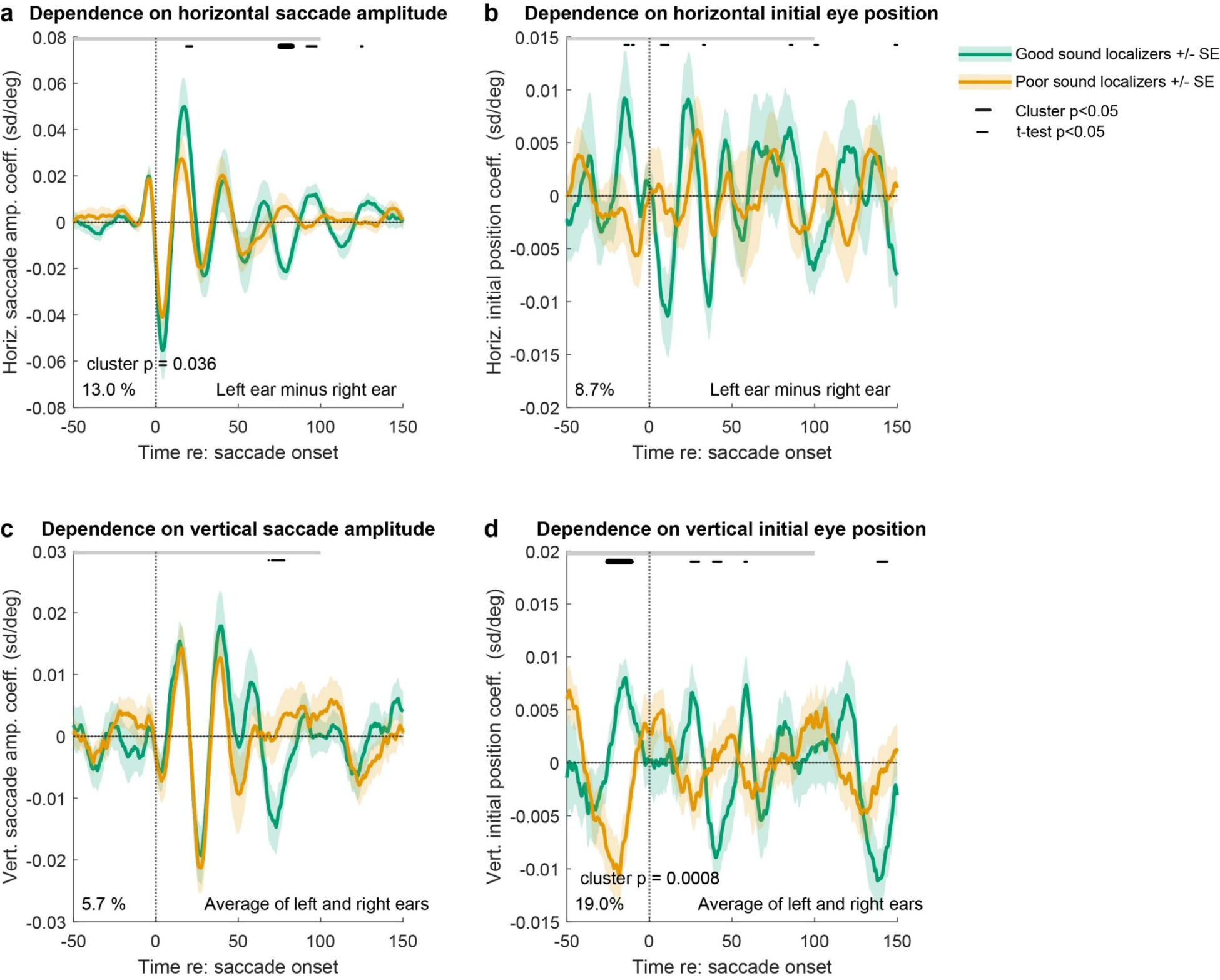
Population average results in “good” vs “poor” sound localizers. Each panel shows the grand average regression coefficient across participants in the “good” (green) or “poor” (orange) groups, with shading indicating standard error of the grand average across the subject population. Individual time points of statistically significant difference between the two groups (two-tailed t-test, p<0.05) are indicated with small black dots at the top of each panel. The total proportion of individually significant time points is shown in the bottom left, for the time period shown with the gray bar (−50 to 100 ms with respect to saccade onset). Periods which passed cluster-based permutation testing (p<0.05) are indicated with larger black dots and the corresponding overall p-values. See main text for full discussion and Supplementary Figures 5 and 6 for left and right ear results presented separately.

Differences between the “good” and “poor” localizers groups can be seen in all four panels, corresponding to the two directions and the saccade amplitude vs initial position regression terms. For horizontal saccade amplitude, the poor localizers (orange) tended to exhibit a smaller signal during the saccade (0-50 ms), and on average the signal disappears sooner than it does for the good localizers. The “good” vs “poor” groups differ at about 13% of the individual time points during a window −50 to 100 ms with respect to saccade onset, exceeding the nominal chance rate of 5%. This was confirmed by cluster-based permutation testing (Maris and Oostenveld 2007) which indicated that the two groups did differ (p<0.05).

For horizontal initial eye position, the signal is overall smaller in the poor localizers and is sometimes out of phase with the signal observed in the good localizers. However, the proportion of individual time points that reach significance by t-test is lower ( 8.7 % −50 to 100 ms time window), and the cluster-based permutation testing does not reach significance (p>0.05).

For vertical saccade amplitude, the signal initially follows a similar pattern across the two groups (e.g for the first 50 ms), but again attenuates earlier in the poor localizers compared to the good localizers. While a brief period of statistical significance of individual time points is evident around 60 ms after saccade onset, the overall proportion of time points significant is close to chance (5.7%) and cluster-based permutation testing does not reach significance (p>0.05).

Finally, the signal related to the initial vertical position of the eyes is quite different among the poor localizers compared to the good ones, exhibiting a phase inversion lasting across several cycles from −25 ms to >50 ms with respect to saccade onsets. Significant differences were observed for 1 9% of the individual time points in the −50 to 1 00 ms time window, and 14.5% overall, and the cluster-based permutation testing reaches an overall significance of p<0.001.

These analyses were repeated subdividing the “poor” localizer group into two subgroups, one for whom the horizontal errors were less than 4.5 degrees but the vertical errors were greater than 5.8 degrees (“moderate”) and another for whom errors in both dimensions were greater than those cutoff values (Supplementary Figure 1). Cluster-based permutation testing between each of the subcategories of “poor” sound localization vs the “good” group confirmed that the disparity was greatest for the initial position of the eyes in the vertical dimension (Supplementary Figure 2) (cluster based permutation testing p<0.05 for both groups) and was also significant for the poorest group’s horizontal saccade dependence compared to the “good” group (cluster based permutation testing p<0.01).

We next visualized the results for each individual “poor” sound localizer, in comparison to the group of “good” sound localizers, focusing on the regression coefficients for horizontal change in eye position and vertical initial eye position, given that these two regression coefficients showed significance in the cluster-based permutation testing. Figures 7 and 8 show each individual “poor” subject, superimposed on the corresponding mean +/- 1 standard deviation across the “good” subjects, with panels organized in descending order of localization accuracy (results for the other two regression coefficients can be found in Supplementary Figures 3 and 4).

**Figure 7.**
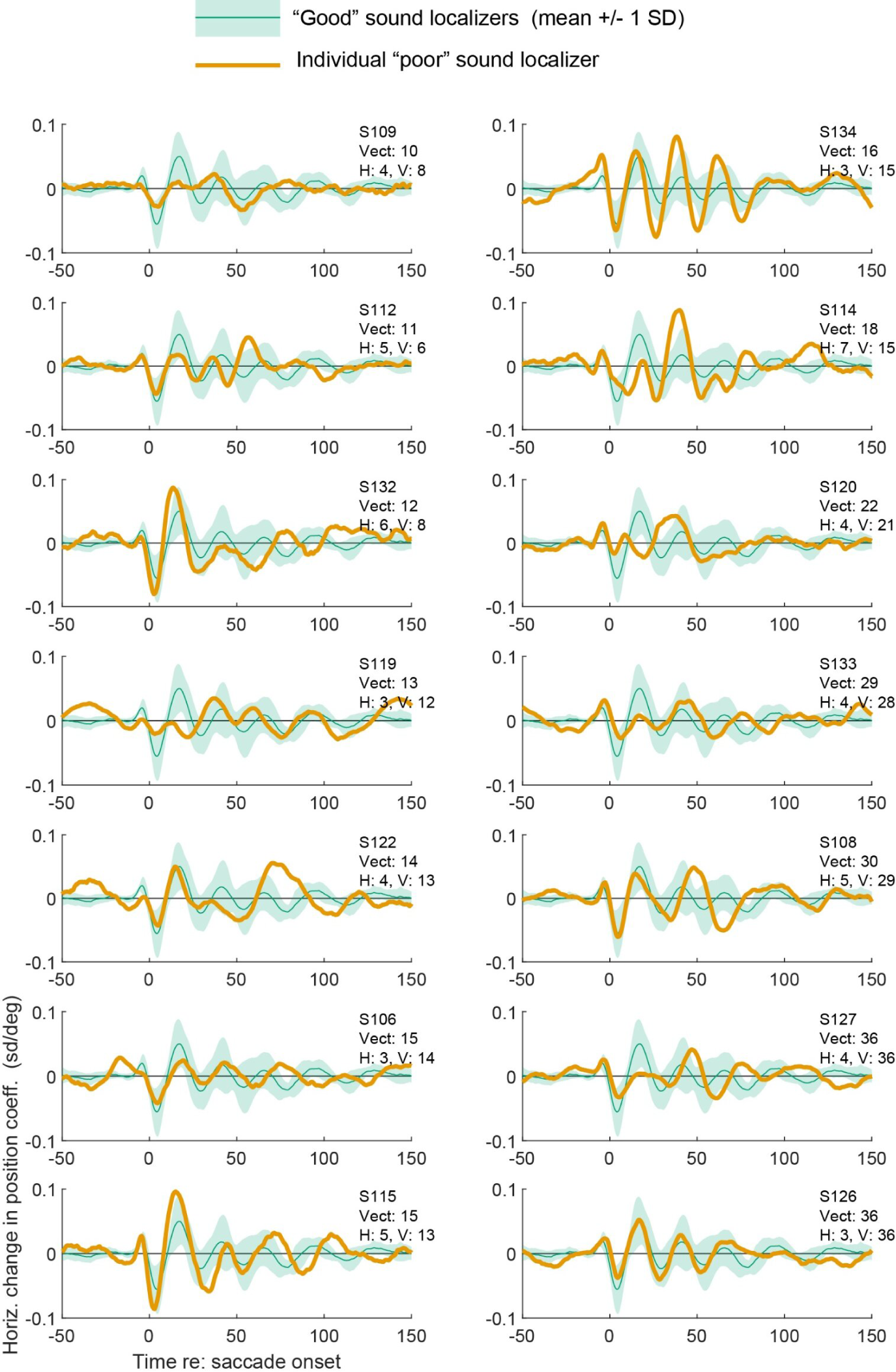
Results of individual subjects from the “poor” localizers group compared to the “good” localizers. Each panel depicts the regression coefficient for the horizontal change in eye position (left ear minus right ear) for one “poor” subject (orange trace) in comparison to the mean +/- 1 standard deviation of the “good” localizer group. Plots are arranged in descending order of localization accuracy; median errors are shown in the top right of each panel.

**Figure 8.**
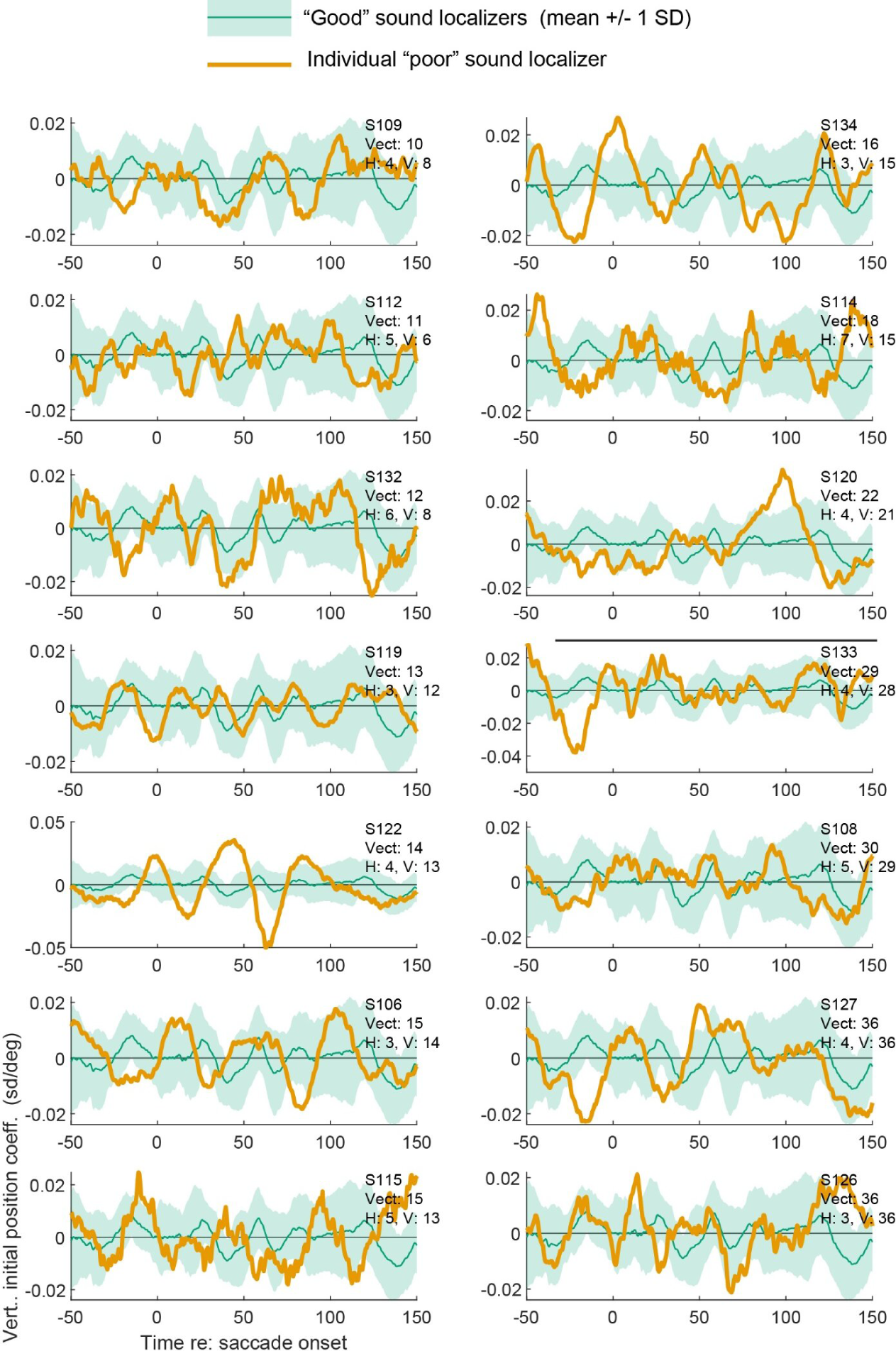
Similar to Figure 7 except the results depict the regression coefficient for the vertical initial position of the eyes (averaged across ears). See Supplementary Figures 3 and 4 for the remaining regression coefficients.

These figures show that there is considerable individual variation across the “poor” localizers, with some showing EMREOs that are on the small side (e.g. Figure 7 S109), others on the large size (e.g. Figure 7 S115, S134). The most common attribute is an unusual shape across time. For the regression coefficient concerning the dependence on the horizontal change in eye position (Figure 7), the most commonly observed atypical pattern involves a late peak/trough that is larger or equal to the peak/trough that occurs in the first 30 ms – this is seen for all but 3 of these 14 participants (the exceptions are S132, S114, and S126). For the coefficient concerning the vertical initial position of the eyes (Figure 8), the pattern is likewise variable across individual participants, but a common attribute is larger oscillations occurring out of phase with those observed among the “good” performers.

## DISCUSSION

Accurate sound localization implies that the brain has performed numerous calculations well: interaural timing differences, interaural level differences, and pinna-based spectral cues have been correctly assessed and assigned to physical locations in the world. Critical to this process is that accurate information about eye position is needed to ascertain where those sound locations lie with respect to the retina/visual scene. Without precise information about eye position, visual calibration of sound location cues across the lifespan would be impossible. We therefore hypothesized that the accuracy of sound localization might be correlated with the properties of eye movement/position signals in the auditory pathway, specifically the EMREO signals found in the ear.

This study presents two main findings addressing this question. First, we found that the accuracy of sound localization via saccades can vary by as much as a factor 12X across individuals – all of whom were deemed to have normal hearing (and normal or corrected-to-normal vision). Second, supporting our hypothesis, the EMREOs differed in subjects with “good” localization abilities vs. those in whom localization was “poor”. Below we consider the implication of the first key finding before turning to the second.

### Individual variation in sound localization as assessed via saccades to sounds

A strength of our study is the use of a naturally-occurring sound localization task: looking at the locations of sounds (a.k.a. auditory-guided saccades) is a normal, everyday behavior. Deficits in this ability have a meaningful real-world impact, potentially making it harder to locate a beeping smoke detector or carry on a conversation with a particular person amidst noisy surroundings. In short, saccade-to-sound tasks preserve a natural link between perception and action and provide an ecologically robust measure of sound localization abilities (Populin 2008). Accordingly, we will first focus our comparison to the previous literature on those that used similar saccade-to sound tasks.

While our study is small (N=23), it is one of the first to be large enough to provide an analysis of the individual variation in saccadic sound localization in normal hearing adults. Previous saccade-to-sound studies have historically relied on scleral eye-coils for eye tracking and thus had small sample sizes (n ranging from 3-6). Such studies have reported vectorial errors ranging from about 1.5 to 8 degrees (Zambarbieri, Schmid et al. 1982, Jay and Sparks 1990, Frens and Van Opstal 1995). Most relevant is a more recent study by Populin (Populin 2008) that noted that subjects (N=9) varied markedly in both accuracy and precision of sound localization in various tasks including gaze shifts and fixation of sound locations. Extracting the most applicable quantitative measures from this study (specifically, the accuracy of fixation of sound locations in frontal space on the horizontal meridian between −29° and 29° - the top section of that study’s Table 3^1^) reveals average vectorial (angular) errors ranged from 2 to 14.5° – i.e a 7-fold range – smaller than what we observed here but confirming that the accuracy of sound localization can vary dramatically across individuals.

### EMREOs in “good” vs “poor” localizers

Numerous aspects of the EMREOs differed between “good” and “poor” localizers. The main implication of this finding is that anomalies in the EMREO may indicate that there are underlying anomalies in the oculomotor information that is incorporated into sound processing and thus the ability to link visual and auditory space. While here we assessed localization abilities as a snapshot in time, over the lifespan visual and oculomotor information are needed to ensure that the underlying cues from which sound location is calculated (interaural timing and level differences as well as spectral cues) remain calibrated.

The import of the specific differences observed in the EMREO is not fully clear. We used a regression method to isolate the aspects of the signal related to the saccade itself (saccade amplitude in the horizontal and vertical dimensions) as well as the initial position of the eyes. Differences were seen in at least two aspects of the EMREO signal – the dependence on the horizontal change in eye position and the dependence on initial position of the eyes in the vertical dimension. The differences observed for the vertical initial position of the eyes were particularly striking, as they involved phase inversions – there were points in time when the EMREO signal in the “good” localizers was going up but the EMREO signal in the “poor” localizers was going down. The saccade amplitude signals largely had a similar shape in both groups, especially during the saccade itself (roughly the first 50 ms after saccade onset), but was smaller in the “poor” group for the horizontal dimension. When results for individual “poor” subjects were scrutinized, a common pattern was that the overall shape of the waveform differed from the “good” subjects, particularly with larger peaks/troughs occurring later in the waveform.

### Comparison to previous EMREO studies

Our study builds on previous attempts to identify a perceptual correlate for EMREOs in normal individuals (Bröhl and Kayser 2023, Sotero Silva, Brohl et al. 2026). Brohl and Kayser (2023) evaluated whether there are any deficits in the ability to detect a sound at the time of an eye movement and Sotero Silva et al (2026) assessed whether the accuracy of judging a lateral shift in sound location was affected by either the direction of a contemporaneous saccade or the timing of the second sound with respect to the EMREO. In the lateralization study, performance was improved when the shift in sound location was the same as the saccade direction, although effects specific to the timing with respect to the EMREO were not apparent. In the sound detection study, no deficits were observed. This latter finding is a reassuring datapoint indicating that EMREOs are not fundamentally deleterious to hearing.

### Spatial hearing – general considerations

As noted above, we used a saccades-to-sounds task partly due to its ecological validity and partly because it is a task that would appear to require the incorporation of information about eye position. Do we expect the results to generalize to other ways of assessing spatial hearing? There are many other ways of testing spatial hearing that do not specifically involve eye movements as a behavioral report (for review, (Freigang, Richter et al. 2015)). There are two possibilities to consider. 1) Eye movements/EMREOs are critical for using visual information to calibrate a generalized representation of sound location in the brain, and the specific form of the behavioral report will not in fact matter, or 2) EMREOs play a role only in sound localization behaviors that directly involve eye movements; tasks such as manual pointing, head turns, left-right judgments, or assessments of minimum audible angle would not show the effects observed here.

Additional studies will be needed to test these competing possibilities. However, many studies that pre-date the discovery of the EMREO do suggest that not everyone compensates for initial eye position absolutely perfectly when localizing sounds. While across studies the effects are variable in magnitude and direction and depend on the paradigm used (Weerts and Thurlow 1971, Lewald and Ehrenstein 1996, Lewald 1997, Lewald 1998, Boucher, Groh et al. 2001, Lewald and Ehrenstein 2001, Metzger, Mullette-Gillman et al. 2004, Lewald and Getzmann 2006, Pavani, Husain et al. 2008, Klingenhoefer and Bremmer 2009, Collins, Heed et al. 2010, Maddox, Pospisil et al. 2014, Krüger, Collins et al. 2016, Willett, Groh et al. 2019) (for review see (Willett, Groh et al. 2019)), nevertheless the parallels with the current work are of interest. Some of the variation in results could be due to chance variation in the subject population when the study is small.

### Limitations and future directions

While our findings indicate that there is a correlation between EMREOs and sound location perception, more work will be needed to understand in what ways different aspects of the EMREOs exert their impact. Our current study was too small to permit evaluation of specific types of anomalies, but with a larger pool of subjects such analyses should become possible.

Another limitation of our current study involves the relationship between features of the EMREO and specifically vertical errors in sound localization. The behavioral sound localization task involved varying horizontal locations, but a single vertical location. Thus, it couldn’t tell us whether subjects could distinguish different vertical locations. Even though vertical location did not vary, a vertical location had to be reported via each saccade, and the vertical errors turned out to be much larger than horizontal errors. This is not surprising because sound localization is known to be less accurate vertically than horizontally, and suggests that assessment of vertical localization abilities may prove to be a more sensitive way to evaluate how well someone can localize sounds. If inaccurate oculomotor cues leading to EMREOs are causally involved in impairing sound localization, vertical accuracy may be the most sensitive to such impairments. A specific correlation between e.g. the horizontal aspect of the EMREO and the horizontal aspect of sound localization is not necessarily expected: an inaccurate sense of eye position in either dimension should lead to errors in sound localization in both dimensions. Put another way, if the auditory system is receiving inaccurate information about eye movements in any direction, that will disrupt calibration of **all** spatial cues, including the spectral cues for elevation.

Insights are also emerging from ongoing studies involving humans with peripheral auditory dysfunction. For example, we recently reported on a human subject who actually hears sounds when she moves her eyes. This person has been diagnosed with tensor tympani myoclonus, and testing in our laboratory showed that she experiences prolonged spasms of her tensor tympani middle ear muscle when she makes large leftward saccades (King, Zhu et al. 2026). The tensor tympani, together with the stapedius middle ear muscle and cochlear outer hair cells, are the likeliest contributors of the internally generated forces that produce the EMREO. This study, then, confirmed that the tensor tympani muscle is indeed a contributor to eye movement-related effects in the ear. This participant’s sound localization abilities were not tested, but this would be a logical next step for investigation.

While we have interpreted our current findings in relationship to physical movements of the eyes, and hypothesized that anomalies in the auditory system’s representation of eye movements could contribute to poor sound localization, this hypothesis doesn’t rule out contributions from other sources. All the subjects in our sample passed basic screening for hearing deficits, but more subtle deficits could have been missed and could have affected both the EMREO and the ability to localize sounds, without the EMREO being causally involved. Similarly, additional variables could affect both EMREOs and sound localization, such as differences in attention. Any possible connection to attention will be of particular interest for future work as there is evidence that attention can itself cause effects in the auditory periphery, potentially independently of eye movements (Smith, Aouad et al. 2012, Srinivasan, Keil et al. 2012, Srinivasan, Keil et al. 2014, Smith and Keil 2015, Dragicevic, Marcenaro et al. 2019, Kohler, Demarchi et al. 2021, Gehmacher, Reisinger et al. 2022). To our knowledge, whether deficits in attentional processing such as ADHD are associated with poor sound localization performance has not yet been studied.

In short, while the hypothesis that the EMREO may be causally involved in computations related to auditory space is plausible on first principles and supported by the correlational evidence presented here and in our previous work, additional steps will be needed to further support this “linking proposition” (Teller 1984). Future work in populations with known types of auditory dysfunction as well as interventional studies in animals will be useful for helping to establish a causal connection between eye movement signals in the ear and auditory perception.

## METHODS

All procedures involving human subjects were approved by Duke University’s Campus Institutional Review Board. Informed consent was obtained from all participants before each testing session, and all subjects received monetary compensation for participation. This study reports data from N=23 human research participants (8 female, 15 male, age range 18-35 yrs, mean age 25.2 years).

Subjects were required to pass audiometry and tympanometry testing to ensure their hearing was within normal limits to be included in the analysis. Specifically, a ll subjects had hearing with air conduction thresholds < 25 dB HL @250, 500, 1000, 2000 and 4000 Hz, determined through pure tone audiometry using a Maico MI-25 screening audiometer and normal middle ear function based on standard 226 Hz probe tone tympanometry (Maico touch Tymp MI-24). Screening audiograms and tympanograms were obtained prior to or on the first day of testing. Middle ear function was assessed with tympanometry on each day of EMREO data collection. Subjects had normal or corrected-to-normal vision. No selection criteria other than an ability to perform the behavioral tasks as noted above were applied. Beyond the N=23 subjects that were included as described above, N=4 subjects were excluded based on excess blinking during task performance.

### Sound localization task

Subjects were seated in a sound attenuation chamber lined with anechoic foam at a distance of 1.25 m from the visual and auditory stimuli display. Subjects were instructed to rest their head on a chin rest with a forehead band to maintain a relatively fixed head position. A video eye tracking system (SR Research Eyelink 1000) provided continuous eye tracking with a sampling rate of 1000 Hz, subsampled by a second data collection computer running Beethoven (Ryklin software) at 500 Hz.

Auditory stimuli consisted of broadband noise at 55 dB SPL and were presented a row of speakers ranging in position from −22° to 22° horizontally at eye level. The speakers were arranged behind a screen of acoustically transparent but visually opaque black cloth. A projector placed above and behind the subjects projected visual stimulus consisting of green dots of approximately 0.5° diameter on the cloth screen, for use as fixation targets or saccade targets as illustrated in Figure 1A.

The task began with an initial fixation for 400-600 ms at one of the three initial fixation positions (−8°,-8 °; 0 °,-8 °; or 8 °,-8 °) before one of the targets in the horizontal row above those fixation positions was presented. Subjects had to maintain fixation in a rectangular window with boundaries within ±5° of the fixation point, for both the horizontal and vertical axes. All trials in which fixation was maintained successfully and the second saccade target was therefore presented were included for analysis, regardless of where the participant looked from that point forward. A 1000 ms grace period was allowed for completing that second saccade, followed by 500 ms of fixation of the second stimulus. The stimuli were presented continuously during that time, for durations that could in principle range from 500-1500 ms depending on saccade reaction time. Visual and auditory trials were randomly interleaved at a 50:50 ratio.

### EMREO recordings and task

Subjects underwent ear canal microphone recordings while performing a visually-guided saccade task in a sound-insulated booth (Acoustic Systems) dedicated to such recordings. The stimuli were presented on a monitor positioned 70 cm away; head position was stabilized using a chin rest. Eye position and ear canal microphone data were recorded while subjects performed the saccade task.

Fixation and target layout is illustrated in Figure 1B. This “Five-Origin-Cross” design is a variant design of previous EMREO experiments that allows assessment of both the effects of initial eye position and saccade amplitude in both horizontal and vertical dimensions in an efficient way. Targets were spaced 9° apart horizontally and 6° apart vertically. Only the fixation position-target location combinations shown were presented, e.g. if the fixation point was leftward, the target would be either to the left or the right but not above or below. The fixation point was illuminated for 750 ms until the target appeared, cueing the subject to make a saccade to that target. No sounds were presented at any time during data collection.

Microphone recordings were obtained in both ear canals simultaneously during all testing sessions using Etymotic Research ER10B+ Low Noise Microphone Systems. A Focusrite Scarlett 2i2 low-latency audio interface provided audio capture and playback at 48 kHz sampling rate through the Etymotic system. Downsampling microphone data from 48 kHz to 2 kHz prior to analysis reduced processing time and allowed the data to remain well above the Nyquist frequency for the range of interest (<100 Hz).

Recordings were made in blocks of ∼125 trials each, with all fixation and target combinations as illustrated in Figure 1B presented pseudorandomly. Each test session consisted of 3 –6 blocks, with breaks offered between blocks. Before each block, the Etymotic ER-2 earphones played a sequence of calibration tones to ensure stability of recording across the testing session (pure tone sweeps from 10 to 1600 Hz: 10 Hz steps for 10–200 Hz, 100 Hz steps for 300:1000 Hz, 200 Hz steps for 1200–1600 Hz). Microphone o utput recorded during tone sweeps was monitored. Any changes (e.g. an earbud dislodging) caused the associated recording block to be discarded. This served to ensure stable microphone function across all testing sessions.

Recording time for each block was approximately five minutes and each test session lasted approximately one and a half hours, including instructions, eye calibrations, microphone assessments, and rest periods.

Each subject’s microphone recording values are Z-scored or expressed in units of standard deviation relative to the variability observed in a 20 ms baseline window 40-60 ms prior to saccade onset, i.e. during a period of steady fixation. The individual microphone samples during this baseline were pooled across all the trials within a recording block and their mean and standard deviation were used to compute the Z-scored values for each of the individual trials in that same block. Data were combined across blocks and sessions.

As for the sound localization sessions, eye tracking information was recorded with the SR Research Eyelink 1000 system, but here the 1 kHz sampling rate was preserved in the saved data. Eye tracking was calibrated at the beginning of each test session using the Eyelink 1000 calibration program and again if the subject moved position or paused for a break in data collection. More details about the justification of methodological decisions and verification of data quality are discussed at length in previously published EMREO studies from our laboratory or collaborators (Gruters, Murphy et al. 2018, King, Lovich et al. 2023, Lovich, King et al. 2023, Lovich, King et al. 2023, Abbasi, King et al. 2025, King and Groh 2026, King, Zhu et al. 2026).

### Statistical analyses

Statistical comparisons of time-series EMREO data were conducted across groups based on their performance on the sound localization task. The basic method followed the procedures laid out in (Maris and Oostenveld 2007), and involved calculating a two-tailed t-test between groups at each individual time point with a criterion of p<0.01, summing the resulting t-statistic over any adjacent time points that passed that criterion (the cluster), and comparing the summed t-statistic(s) to a shuffled distribution to generate a p-value for each cluster. If at least one cluster exhibited a p-value < 0.05, the groups were considered to show a significant difference.

## Acknowledgments

This work was supported by NIH grant DC020363 to JMG and CDK. We thank Stephanie N. Lovich for thoughtful comments on the analysis and manuscript.

## SUPPLEMENTARY FIGURES

**Supplementary Figure 1.**
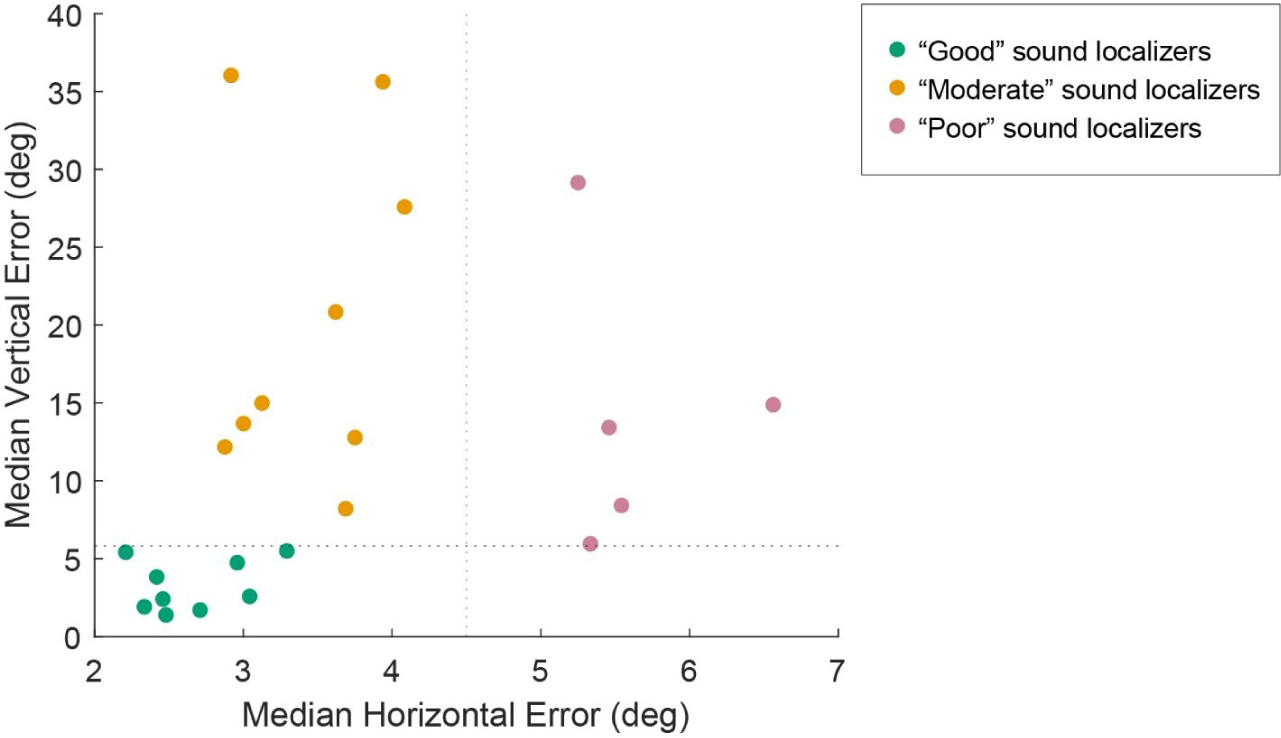
Criteria for subdividing “poor” sound localizers into two subgroups, for the analysis presented in Supplementary Figure 2.

**Supplementary Figure 2.**
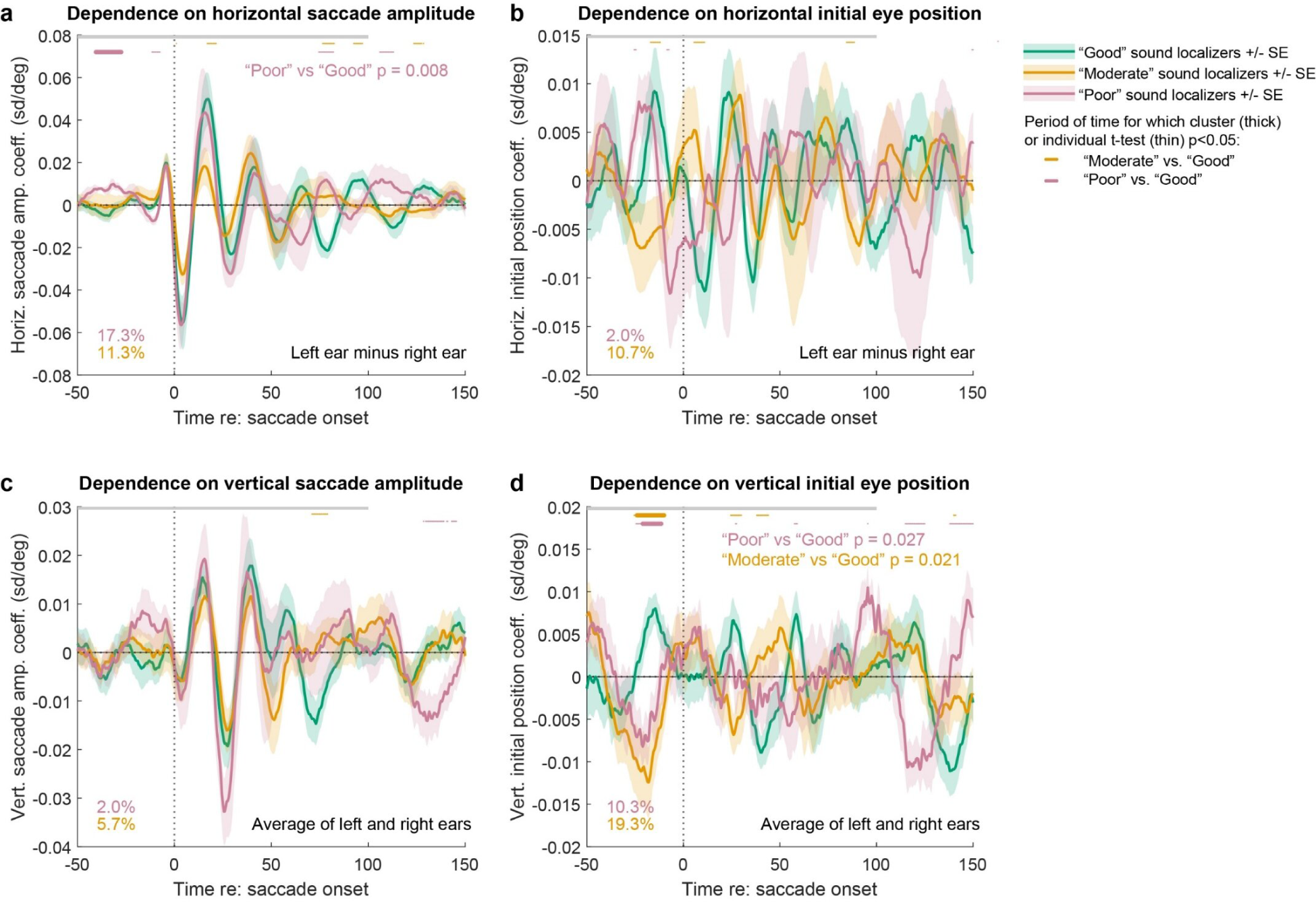
Population average results for “good” vs “moderate” vs “poor” sound localizers. Each panel shows the grand average regression coefficient across participants in the “good” (green) or “moderate” (orange) or “poor” groups (pink), with shading indicating standard error of the grand average across the subject population. Periods of statistically significant difference (two-tailed t-test, p<0.05) are indicated with orange and pink dots at the top of each panel, and the total proportion of significant time points is shown in the bottom left, for the time period shown (−50 to 100 ms with respect to saccade onset, gray bar). Cluster based permutation testing results are as indicated.

**Supplementary Figure 3.**
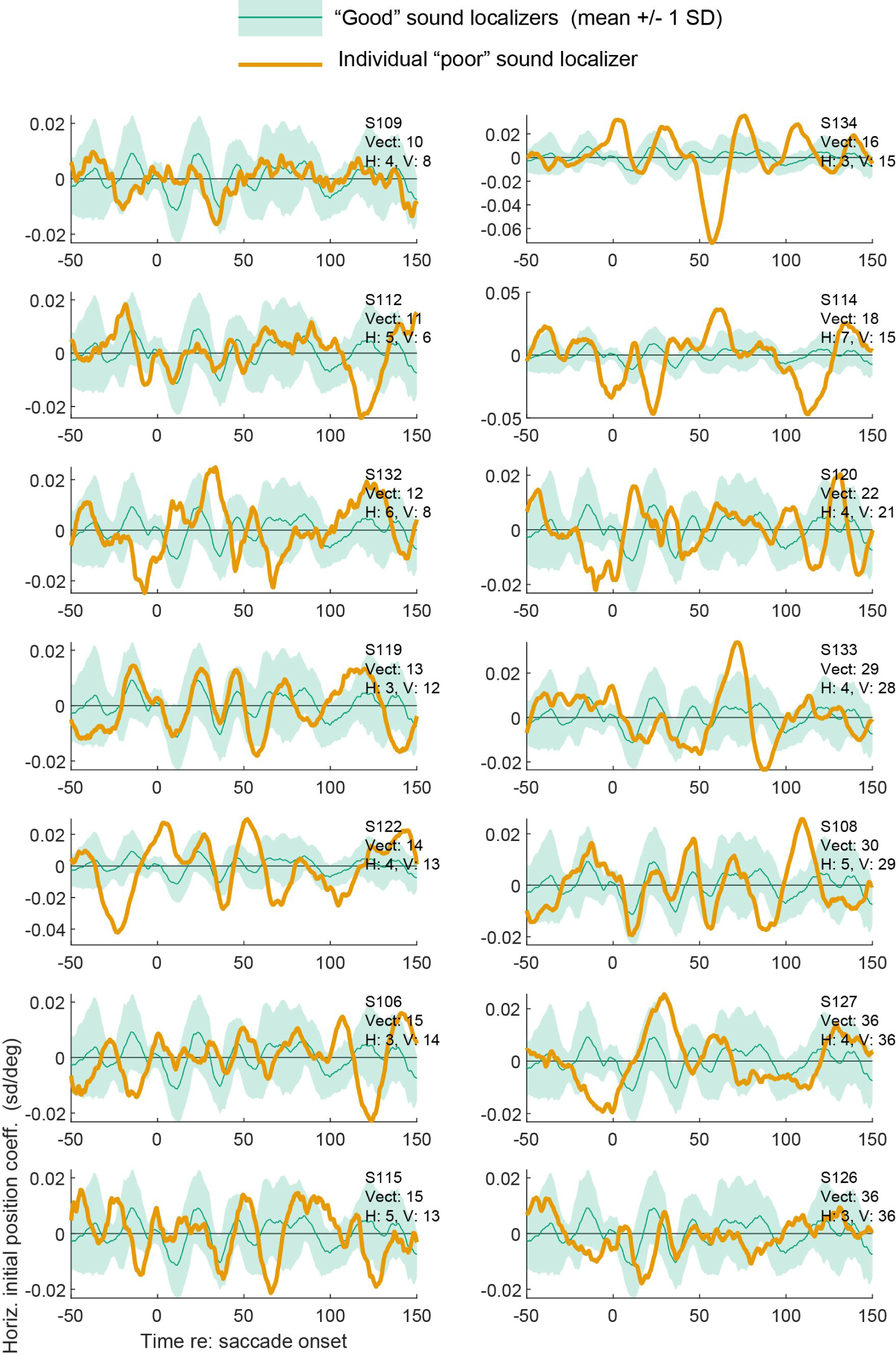
Similar to Figure 7 except the results depict the regression coefficient for the horizontal initial position of the eyes (left ear minus right ear).

**Supplementary Figure 4.**
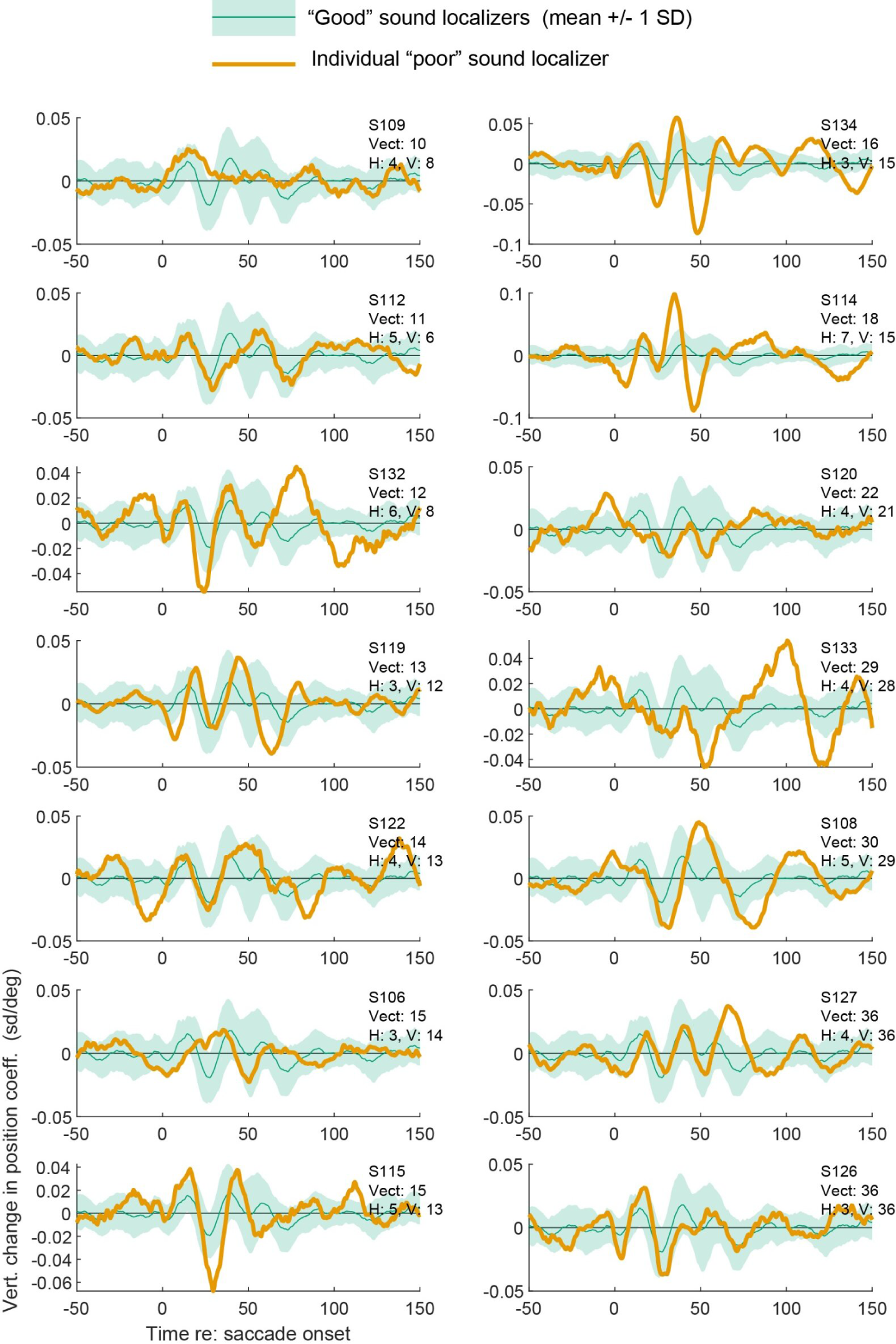
Similar to Figure 8 except the results depict the regression coefficient for the vertical change in position of the eyes (averaged across ears).

**Supplementary Figure 5.**
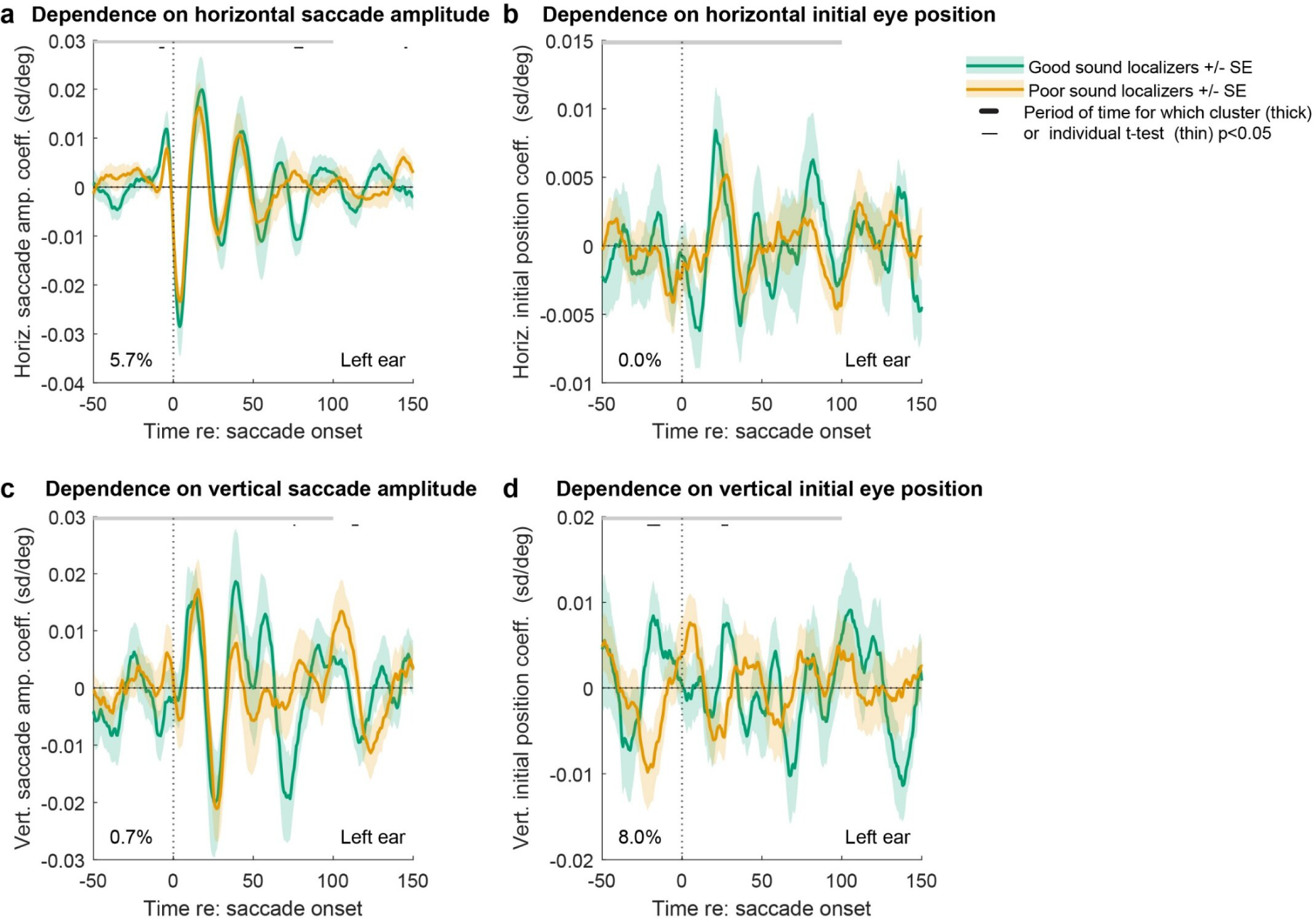
Similar to Figure 6 except results are for left ear. Cluster-based permutation testing did not reveal significance (p>0.05).

**Supplementary Figure 6.**
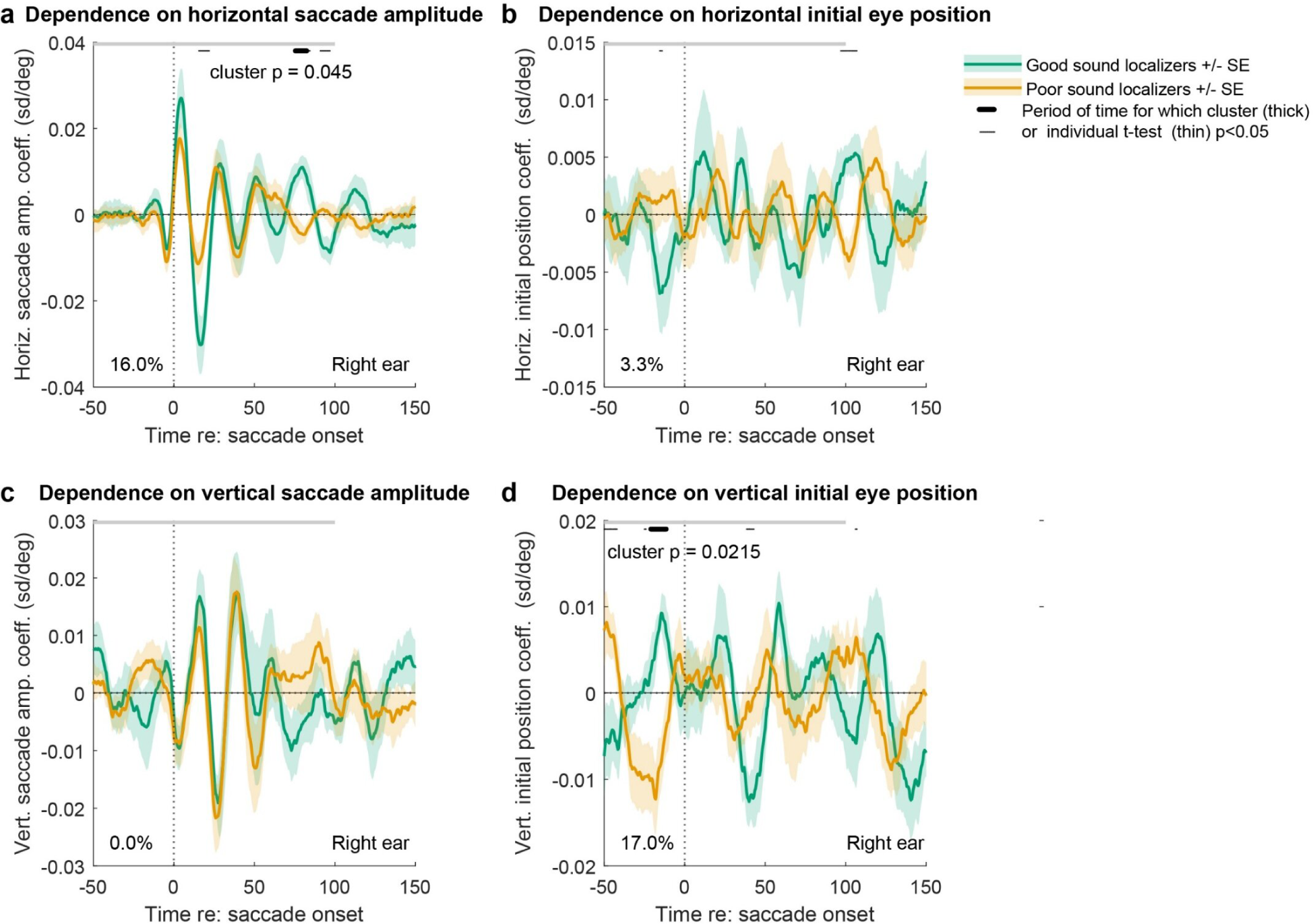
Similar to Figure 6 except results are for right ear. Cluster-based permutation testing revealed significance for the dependence on horizontal saccade amplitude and vertical initial eye position (p<0.05).

## Footnotes

1 See repository of data and code for re-analysis of Populin 2008’s Table 3.

